# ECMME: an atlas of selection pressures on the mammalian extracellular matrix reveals contrasting evolutionary dynamics

**DOI:** 10.64898/2026.06.08.730821

**Authors:** Petar B. Petrov, Ayomide Oshinjo, Juha Röning, Valerio Izzi

## Abstract

The extracellular matrix (ECM) is a fundamental metazoan innovation that provides structural support and regulatory cues essential for multicellular life. While core matrisome components are subject to strong functional constraints, their evolutionary dynamics at the molecular level remain incompletely characterized. Here, we present a comprehensive per-residue analysis of selection pressures across 272 human core matrisome proteins using high-quality orthologous sequences from up to 228 placental mammal species. We developed an automated pipeline integrating ortholog identification, codon-aware alignments, and site-specific selection analyses with the MEME and FUBAR methods from the HyPhy suite. Results reveal pervasive strong purifying selection across the matrisome, consistent with its structural and functional indispensability. This is accompanied by episodic positive selection and rarer pervasive positive selection, with collagens exhibiting significantly elevated episodic positive selection compared to glycoproteins and proteoglycans.

To facilitate community access, we developed ECMME (ECM Molecular Evolution) browser, an intuitive open-access web resource that visualizes selection metrics plotted directly onto protein topologies. ECMME allows researchers to seamlessly browse and investigate the data, providing a powerful framework for interpreting functional sites. It is available online and requires no local installation or set-up (https://izzilab-ecmme.share.connect.posit.cloud/).

## Introduction

Computational phylogenetic studies provide a powerful framework to investigate the evolutionary history of genes and consequently the proteins they encode. Evolutionary selection analyses estimate the relative non-synonymous (dN) and synonymous (dS) substitution rates across the coding sequences (CDS) of a gene’s orthologues, allowing inference of the type and strength of natural selection. The predominant form of selection is negative (purifying), which eliminates deleterious mutations, thus maintaining functional integrity [1]. Housekeeping and essential genes typically evolve under strong purifying selection, resulting in high sequence conservation evident across species [2]. In contrast, the much more rare positive (Darwinian) selection favors advantageous mutations and promotes adaptation. It has been frequently detected in genes involved in immunity, reproduction, sensory perception, and other systems where rapid molecular innovation confers fitness advantages [3].

The extracellular matrix (ECM) is a three-dimensional protein scaffold that surrounds nearly every cell in metazoan tissues, providing structural support together with biochemical and mechanical cues that regulate cell adhesion, migration, proliferation, differentiation, and survival [4,5]. The ECM proper – formally, the core matrisome – comprises approximately 300 mammalian genes encoding collagens, laminins, fibronectins, elastin, and proteoglycans [6], whose functions are so fundamentally linked to life that its dysregulation underlies or accompanies virtually any disease, including fibrosis, tumor progression, congenital connective-tissue disorders, and ageing [7,8]. From an evolutionary perspective, the ECM is an ancient innovation: collagen- and laminin-like networks predate the bilaterian radiation and accompanied the transition to multicellularity [9]. Yet the ECM proper is far from evolutionarily static. Across mammals, its genes display a striking duality – deep purifying selection on canonical structural elements, such as the Gly-X-Y triple helix of fibrillar collagens, alongside rapid diversification of regulatory domains, alternatively spliced exons, and matricellular signalling motifs [9,10]. This balance of constraint and innovation is fundamental to the existence of ECM itself. However, while substantial progress has been made in characterizing ECM proteins, their selective pressures at the molecular level remain largely underexplored, leaving their patterns and functional impact enigmatic.

Here, we performed a comprehensive and detailed phylogenetic analysis of the per-residue evolutionary pressure in 272 core matrisome proteins. Automation becomes essential for large-scale data analysis, making the design of a tailored pipeline a fundamental part of the work. By taking advantage of the R language and BASH scripting, we achieved a high level of systemization for the hereby presented work. This yielded a detailed dataset, that integrates quality-filtered placental mammals ortholog sequences, codon-aware alignments and *site-* and *branch-site* tests for selection, implemented in HyPhy [11,12]. Containing the orthologues to level *Placentalia* mitigates the risk of sequences misalignment in the subsequent steps of the study, thereby reducing false-positive results. We developed ECMME -- the ECM Molecular Evolution Browser -- an intuitive online interface for the interrogation of our database, enabling researchers to effortlessly search, explore, and analyze proteins of interest, with evolutionary insights directly visualized onto the protein topology. For every residue in the 272 core matrisome proteins, the resource provides calibrated metrics of purifying, neutral, and positive selection, together with all underlying alignments. Ortholog assignments, processing scripts, and quality-control reports required for full reproducibility and further analysis are also made publicly available. We anticipate that the dataset will support mapping selection onto protein structures and binding interfaces, prioritising functional residues for mutagenesis, interpreting disease-associated variants, and reconstructing the evolutionary trajectories of matrix assembly and remodelling across mammals.

## Results

In this study, we curated a high-confidence set of 274 human extracellular matrix (ECM) components corresponding to the “core matrisome” as defined by Naba et al, 2012 [13] and available through the Matrisome Project portal (https://naba.lab.uic.edu/the-matrisome-project). Building on this reference, we implemented our comprehensive and scalable pipeline to identify orthologous sequences across placental mammals for each component (Figure 1; see Materials and Methods and Code Availability). For consistency in downstream analyses and visualization, we adopted the “canonical” human isoforms as defined in UniProt for 266 proteins (“Data availability, Additional File 1”). In the remaining eight cases where no direct correspondence was available in NCBI RefSeq, we instead selected the canonical RefSeq isoform. Protein-level annotations, including domain architecture, short functional motifs (SFMs), and post-translational modifications (PTMs), were systematically retrieved from the corresponding database, ensuring harmonized feature mapping. We searched NCBI for homologous sequences for the 274 core ECM members, by their Entrez gene IDs at level *placentalia*. Following stringent sequence quality control and taxonomic validation using TimeTree [14], we obtained orthologous sequences spanning 228 species (Figure 2, Supplementary Table S1, Supplementary Figure S1). The tree topology featured seven distinct clades, five major and two minor, depending on their number of species. The five major clades (25-54 species each), covered Orders: Primates, Rodentia, Carnivora, Artiodactyla and Chiroptera. Notably, species representation varied across individual genes, precluding the definition of a uniform species set of consistent size across all ECM components, the median species coverage per gene being 186 (“Data availability”, Additional File 2). All candidate ortholog sequences were subjected to reciprocal BLAST against the corresponding human reference, then filtered and ranked to enable the selection of a single, high-confidence isoform per species for each ECM component. Downstream analyses therefore encompass 272 ECM proteins, excluding one pseudogene and one protein for which only the human sequence satisfied all quality thresholds.

**Figure 1.**
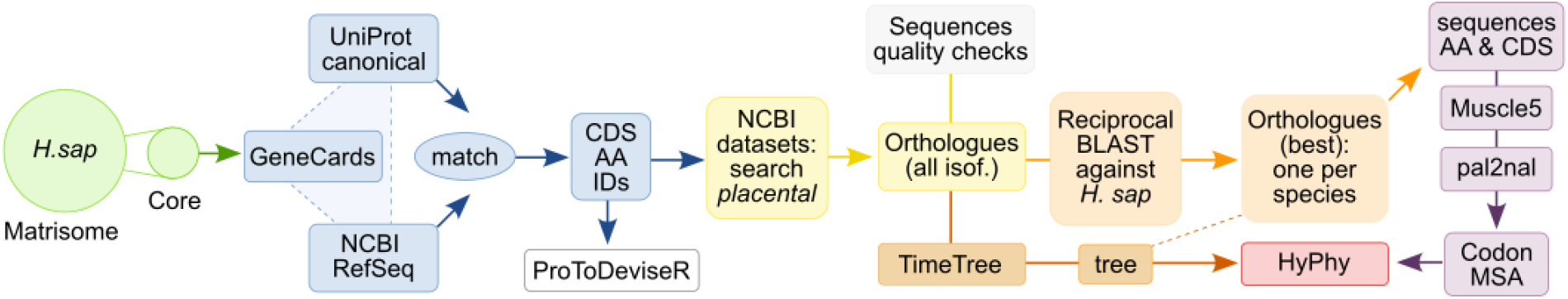
Orthologues identification and evolutionary analysis pipeline. The UniProt IDs of the core members of the Matrisome from *H. sapiens* (green) are cross-referenced against GeneCards, UniProt and NCBI RefSeq, yielding a list of updated IDs and sequences (blue). The UniProt IDs are used to generate protein topology schemes in JSON format by ProToDeviseR (white). Orthologues are then retrieved by NCBI Datasets at level *Placentalia* (yellow), followed by sequence quality checks and filtering. Reciprocal BLAST searches against the human reference are performed to select the best orthologue per species (orange). Amino acid sequences for the orthologues of each Matrisome core member are aligned with MUSCLE5 and the corresponding CDS are converted to codon alignments using pal2nal (purple). The codon alignments are used together with a species tree (brown), obtained from TimeTree (trimmed accordingly, to represent only the species available in the CDS) as input for HyPhy evolutionary analyses by methods MEME and FUBAR (red).

**Figure 2.**
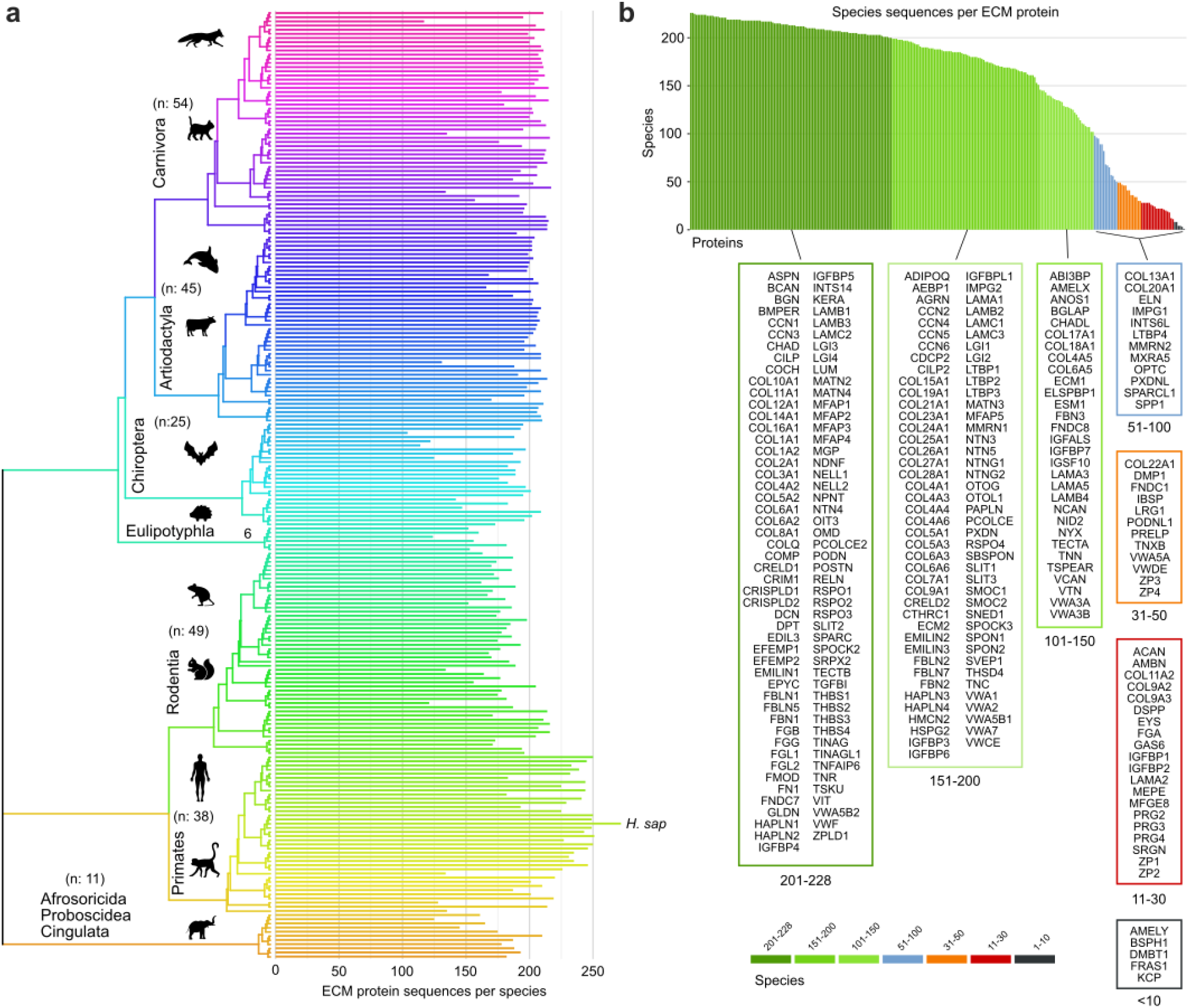
Species and orthologues. A) Species included in the analysis. A total of 228 species are shown using a rainbow colour scheme, ordered according to their position in the unrooted phylogenetic tree. The tree topology highlights seven major clades, with the primary taxonomic orders representing the majority of species in each clade indicated. Species counts per clade are shown in brackets. The number of ECM protein sequences retained per species after filtering is displayed as absolute values, with Homo sapiens highlighted. **B) Number of species per ECM protein**. Molecules are grouped according to the number of species in which an orthologous sequence was successfully identified following screening.

To interrogate the evolutionary forces shaping the core matrisome, we constructed codon-level alignments of orthologous sequences and mapped them onto a global species phylogeny. Selection pressures were then quantified using the MEME and FUBAR frameworks implemented in the HyPhy suite. Across the ECM components analyzed, the prevailing signal was one of strong purifying (negative) selection (Figure 3A), consistent with the structural and functional indispensability of the extracellular matrix [x]. This dominant constraint, however, was accompanied by signatures of episodic and, more rarely, pervasive positive selection within the same molecule, pointing to localized adaptive fine-tuning.To glimpse into the possible structural ties of molecular evolution of the ECM, we next examined how these evolutionary regimes distribute across the principal matrisome divisions. The 272 ECM proteins partition into three non-overlapping classes of ECM glycoproteins, collagens, and proteoglycans, with glycoproteins comprising the largest fraction (Figure 3B). Strikingly, collagens exhibit a significantly elevated burden of episodic positive selection relative to the other divisions (Figure 3C), suggesting that even the *allegedly* most structurally-constrained ECM class retains pockets of adaptive evolutionary flexibility, potentially linked to species-specific biochemical, biomechanical or functional demands. In contrast, no statistically significant differences were observed for pervasive positive or negative selection across the three groups, reinforcing the notion of a shared, dominant baseline of evolutionary stability across the ECM.

**Figure 3.**
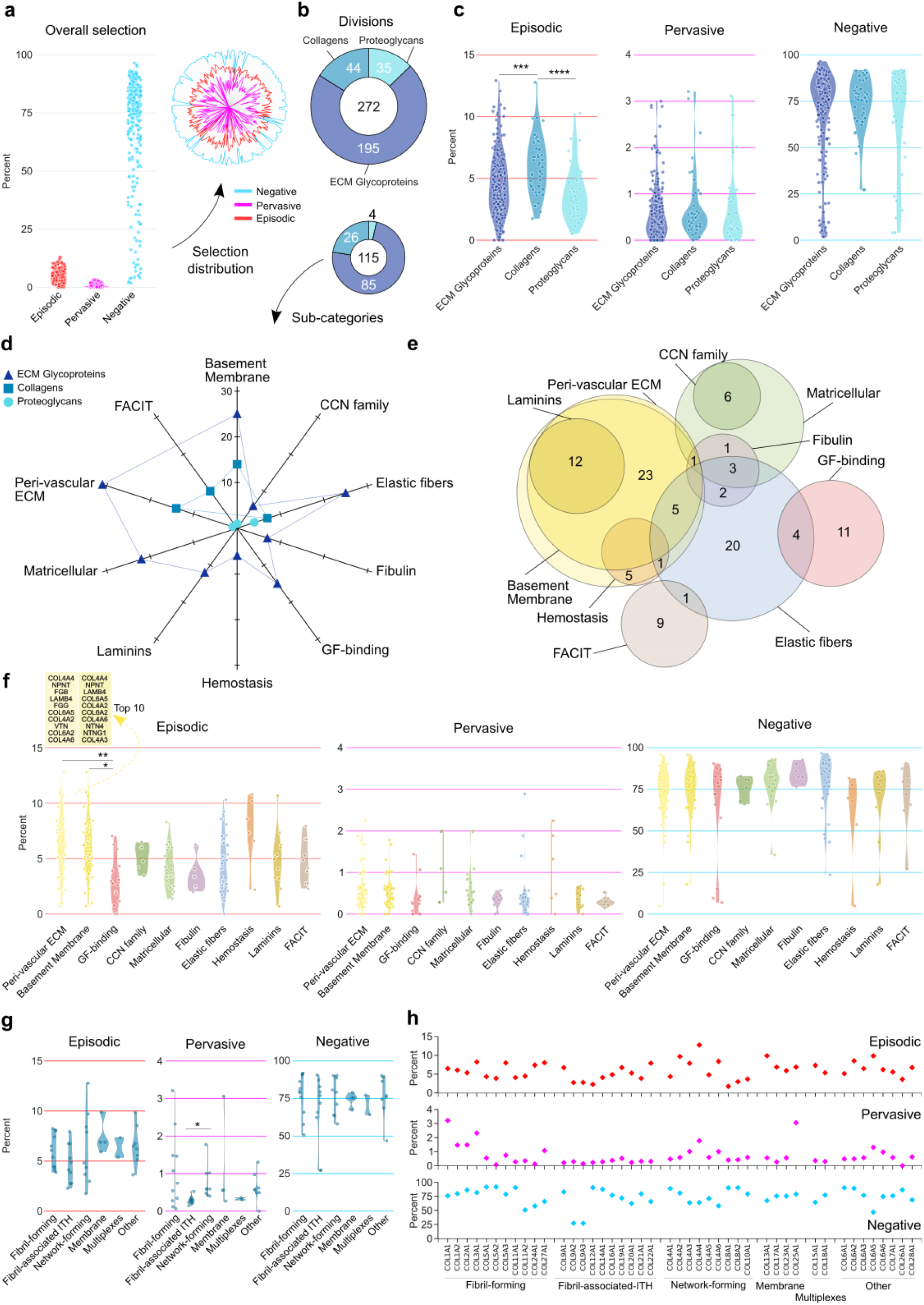
Evolutionary selection across the dataset. **A) Overall selection. Left:** Episodic, pervasive positive and negative selection on molecules, as percentage of sites. **Right:** Distribution of selection across the molecules, as an overlapping radar plot. Scale is logarithmic (center - edge, 0 - 100) for better readability. **B) Classifications of ECM proteins**. The 272 molecules are split into 3 non-overlapping major divisions (up), while a subset of 115 proteins (down) is further annotated into 10 overlapping sub-categories (D, see below). **C) Selection on major divisions**. Episodic, pervasive positive and negative selection on ECM glycoproteins, collagens and proteoglycans. Statistically significant difference was observed for the episodic selection between ECM glycoproteins and collagens, by pairwise Wilcox test, with Holm corrected p-values (*** p < 0.001, **** p < 0.0001). **D) Sub-categories**. The major divisions allocated to the 10 sub-categories. **E) Overlap between the sub-categories**. Protein counts are shown as absolute numbers. **F) Selection across sub-categories**. Episodic, pervasive positive and negative selection on the 10 sub-categories. Statistically significant difference was observed for the episodic selection between Peri-vascular ECM / Basement membrane (largely overlapping sub-categories) and Growth-factor (GF) binding by pairwise Wilcox test, with Holm corrected p-values (* p < 0.05, ** p < 0.01). **G) Selection on collagen sub-categories**. Statistically signifficant difference was found between the pervasive selection of Fibril associated with interrupted triple helices (ITH) and Network-forming collagens, by pairwise Wilcox text, with Holm corrected p-values (* p < 0.05). **H) Selection on individual collagens**.

We obtained comparable results at finer functional resolution, when we further stratified a subset of 115 ECM components into ten subcategories (Figure 3B, lower panel; Figure 3D). These subsets are dominated by ECM glycoproteins and collagens, with only a minor contribution from proteoglycans. Notably, the subcategories are not mutually exclusive (Figure 3E), with the greatest overlap observed among peri-vascular ECM, basement membrane constituents, and laminins -- all tied by architectural and functional interdependence. Within this refined framework, selection patterns largely follow the patterns precedingly described, with peri-vascular ECM and basement membrane groups displaying an enrichment of episodic positive selection relative to growth factor (GF)–binding proteins (Figure 3F). The “top 10” molecules, ranked by episodic selection, from the two largely overlapping subcategories constitute a relatively equal numbers of collagens and ECM glycoproteins, thus the elevated episodic selection is not solely due to the presence of collagens. Hence, the structural ECM compartments may be preferential substrates for adaptive change against a broadly conserved evolutionary landscape, especially in comparison to GF-binding proteins

Then we focused on the selection across collagens, the major division that displayed highes rates of episodic selection. Intriguingly, we detected statistically significant difference in the pervasive episodic selection between Fibril-associated collagens with interrupted triple helices and Network forming.

### Evolutionary pressure at the single-molecule scale

As molecular examples, we highlight top-ranking proteins from each selection regime: episodic/diversifying, pervasive positive, and negative (purifying) selection. For improved visualization, these molecules were chosen for their well annotated distinct domains, short functional motifs, and post-translational modification sites (PTMs): EMID1 (EMI domain-containing protein 1), COL1A1 (Collagen alpha-1(I) chain), and NTN1 (Netrin-1), respectively.

#### Overall Protein Topology

Detailed protein feature annotations were retrieved automatically from UniProt using ProToDeviseR — the same tool employed to generate the graphical protein topologies displayed in ECMME. The three proteins exhibit distinct domain architectures: EMID1 contains an EMI domain and a collagen-like (Col-like) domain; COL1A1 features VWFC, THR, and FCNC1 domains; and NTN1 is characterized by Laminin, L-EGF-L, and TNC domains (see Figure 4A legend for full details and abbreviations). In addition, EMID1 and COL1A1 harbor long disordered regions, low-complexity segments, and proline-rich or polar/charged residue stretches. All three molecules also display patterns of glycosylation, disulfide bridges, and other PTMs. Notably, COL1A1 contains numerous hydroxyprolines within the THR and two intermolecular disulfide bridges at the FCNC1 domain, whereas NTN1 shows the highest proportion of intramolecular disulfide bonds.

**Figure 4.**
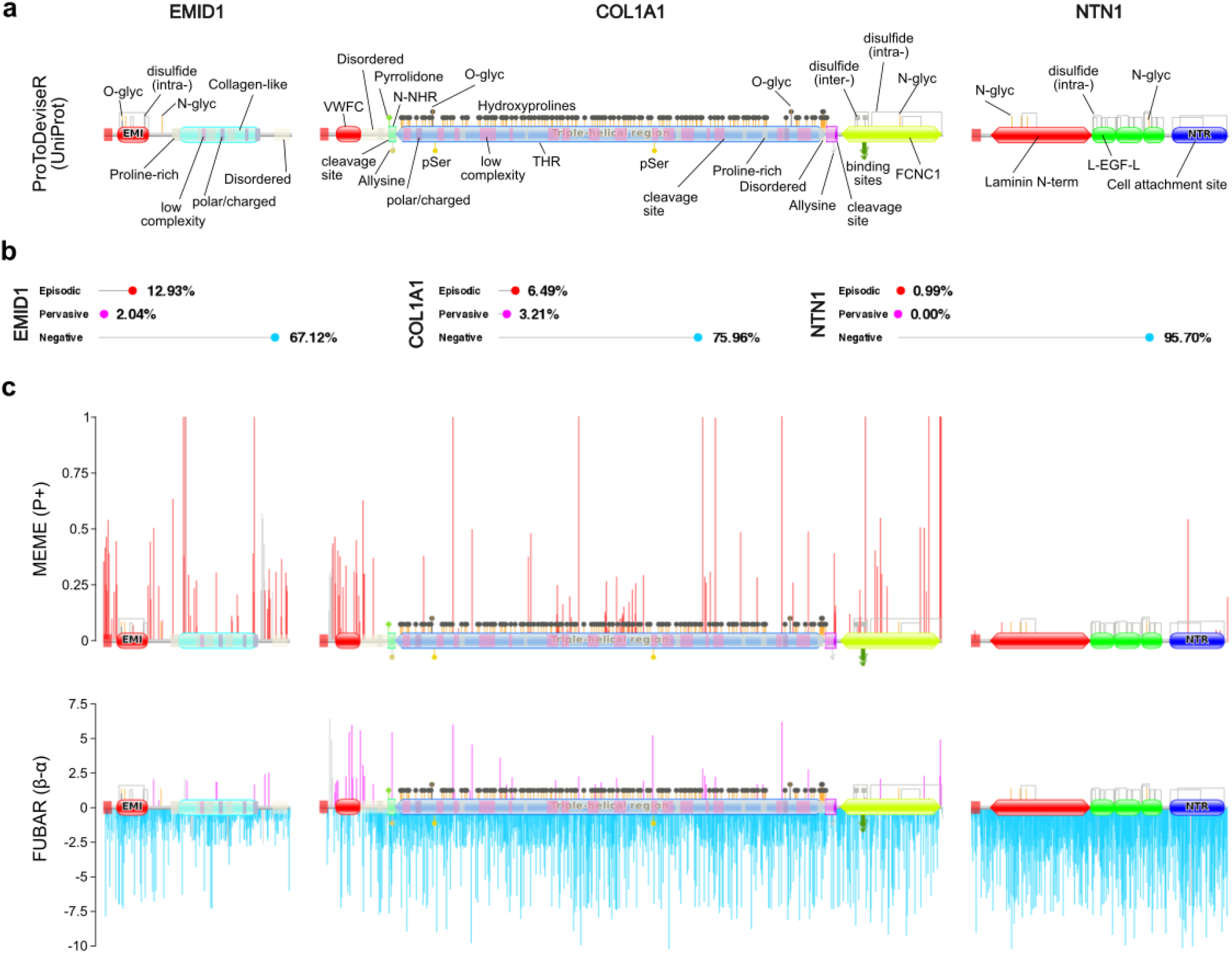
Evolutionary selection on EMID1, COL1A1 and NTN1. **A) Protein sequences topology. Left:** the canonical isoform (Q96A84-1) of EMID1 is a 441-aa long protein, that features a signal peptide, an EMI domain and a Collagen-like domain. A large disordered region starts upstream from the Collagen-like domain, completely overlapping with it. The Collagen-like domain also contains a low-complexity motif and interspaced proline-rich and polar/charged amino acid motifs. A smaller disordered region resides at the very C-terminus of the protein. Three intramolecular disulfide bridges are found within the EMI domain, as well as an O- and an N-glycosylation site. Another N-glycosylation site resides in the space between the EMI and Collagen-like domains. **Middle:** The canonical isoform (P02452) of COL1A1 is 1464-aa long and starts with a signal peptide, followed by a VWFC (Von Willebrand factor, type C) domain, a N-NHR (N-terminal non helical region), a THR (triple-helical region), a C-NHR (C-terminal non-helical region) and a C-terminal FCNC1 (Fibrillar collagen non-collagenous 1) domain. The section between the VWFC domain and the near end of C-NHR is annotated as disordered, containing proline-rich, charged/polar and low-complexity motifs, most of which are found within the nested THR. The N-NHR is preceded by a cleavage site, and contains a pyrrolidone carboxylic acid modified residue, an allysine and a serine phosphorylation site. The THR is abundant in hydroxyproline sites, and also accommodates two serine phosphorylation sites, two O-glycosylation residues and a cleavage site. The C-NHR contains an allysine and the third cleavage site of the molecule, situated at its very end. The FCNC1 domain features three intra-molecular and two inter-chain disulfide bridges, five binding sites and an N-glycosylation site. **Right:** The canonical isoform of NTN1 (O95631) is 604-aa long, beginning with a signal peptide, followed by a LNT (Laminin N-terminal) domain, three L-EGF-L (Laminin Epidermal Growth Factor like) domains and a NTR (Netrin) domain at the C-terminus. A single short functional motif is annotated for the molecule, a cell attachment site situated within the NTR. Total of 15 intramolecular disulfide bridges are listed for NTN1: 1 in the LNT domain, 12 over the L-EGF-L domains and 2 in NTR. The LNT domain bears 3 N-glycosylation sites, while a fourth one is found in the last L-EGF-L domain. **B) Fractions of sites under episodic, pervasive and negative selection. C) Evolutionary selection. Left:** In EMID1, episodic selection was abundant at the signal peptide, around the EMI domain and towards the C-terminus, while patches of selected sites were found in the beginning of the Collagen-like domain. Pervasive positive selection was detected between the domains, withing the Collagen-like domain, and towards the C-terminus. Negative selection appeared strongest in the N-terminal region and following the EMI domain, as well as at a proline-rich motif within the Collagen-like domain. **Middle:** In COL1A1, sites under pervasive positive selection were distributed mainly around and within the VWFC domain and to a lesser extent throughout the THR, the FCNC1 domain and the C-terminus. The protein also features abundant episodic selection, following a similar distribution, often overlapping with the pervasive selection. Negative selection appeared somewhat less-abundant at the N-terminus, where the VWFC domain is situated. **Right:** In NTN1, negative selection was particularly high in the NTR domain. The small fraction of sites under episodic diversification, were predominantly within the NTR domain, while none were found to be under pervasive positive selection.

#### Overall Evolutionary Selection

We calculated the overall selection pressure on each protein as the percentage of sites under selection relative to the protein’s total length (amino acid count; Figure 4B). EMID1 stood out with the highest prevalence of episodic (diversifying) selection in our dataset (Figure 4C, left), particularly concentrated at the borders of the Col-like domain and inter-domain regions. Sites under pervasive positive selection were far less common but still followed a similar distribution. In contrast, COL1A1 showed the highest fraction of sites under pervasive positive selection (Figure 4C, middle), especially within the VWFC domain, with additional sites scattered across the rest of the molecule. Episodic selection in COL1A1 largely followed the same distribution and somewhat flanked the pervasive positive selection at VWFC. Negative (purifying) selection was widespread across all three proteins, featuring strongest levels in NTN1 (Figure 4C, right), particularly within the NTR domain. Somewhat contracting, a few sites within this same NTR domain also showed evidence of episodic selection.

#### Functional Motifs Prediction and Selection

To complement the existing UniProt annotations, we predicted short functional motifs (SFMs) in EMID1, COL1A1, and NTN1 using the ELM resource online. In total, we identified 11 different types of SFMs across the three proteins (Figure 5A), featuring various motifs for cleavage, amidation, glycosaminoglycan attachment, N-glycosylation, fucose attachment, as well as, collagen-specific binding, and integrin RGD-type binding. Five of these motifs for cleavage sites, amidation, N-glycosylation, and glycosaminoglycan attachment were shared among all three proteins (Figure 5C). Examining the average selection signals (P+ or β-α from MEME and FUBAR) across these motifs (Figure 5B), revealed that all SFM types were predominantly under negative selection. Motifs with distinct episodic selection were those for glycosaminoglycan attachment and several cleavage sites. Notably, despite NTN1 exhibiting the strongest overall negative selection, one of its cleavage motifs showed signs of episodic diversefication. EMID1, on the other hand, was unique in harboring a cleavage motif that contained sites under pervasive positive selection.

**Figure 5.**
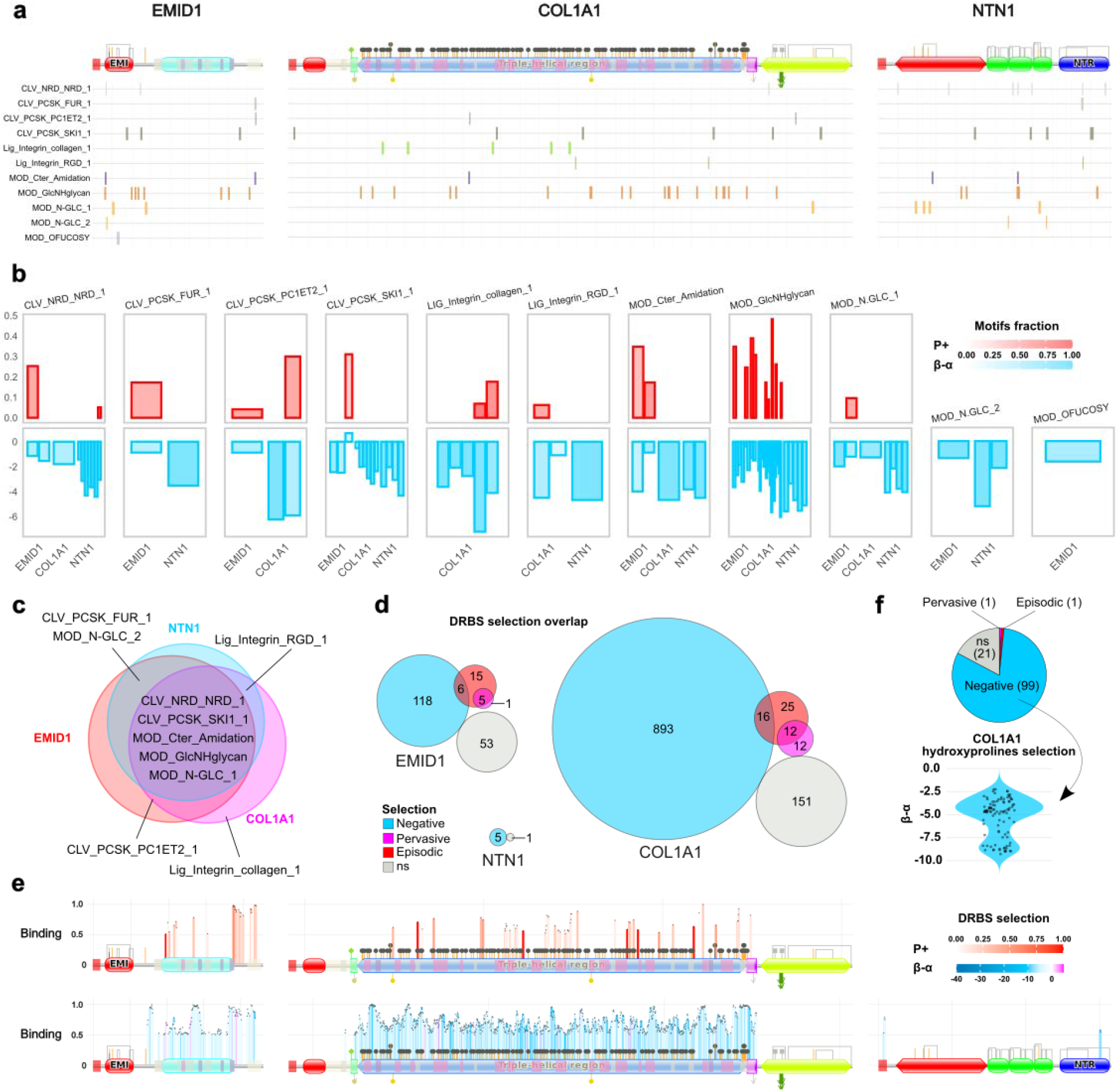
Patterns of evolutionary selection on motifs and disordered region binding sites (DRBS). **A) Predicted short functional motifs**. Total of 11 types of short functional motifs (SFMs) were identified for the three molecules altogether. The SFMs feature cleavage sites (CLV_NRD_NRD_1, CLV_PCSK_FUR_1, CLV_PCSK_PC1ET2_1, CLV_PCSK_SKI1_1), amidation (MOD_Cter_Amidation), Glycosaminoglycans attachment sites (MOD_GlcNHglycan), N-glycosylation (MOD_N-GLC_1, MOD_N-GLC_2), fucose residue attachment (MOD_OFUCOSY), collagen-specific binding motif (LIG_Integrin_collagen_1) and an Integrin RGD-type binding site (LIG_Integrin_RGD_1). **B) Selection on motifs**. Mean values for the selection, as inferred by MEME and FUBAR, within each motif are shown, by averaging either P+ or β-α, estimated for the motif sites. Colour intensity corresponds to the fraction of amino acids per motif that are under selection, discarding the ones that did not pass the statistical significance tests of the methods. **C) Overlap between the predicted SFMs. D) Overall selection DRBS**. Sites per molecule are shown as absolute numbers. **E) Selection on DRBS across the molecules. Left:** In EMID1, DRBS with high binding scores (>0.8) were inferred to be under episodic selection within the post Collagen-like domain area was. Analogously, negative selection was detected for high binding scores DRBS preceding, superseding and in the middle of the Collagen-like domain itself. **Middle:** In COL1A1, DRBS under negative selection show 7 “peaks” of particularly high binding scores, across the THR and its borders. Most of the DRBS under episodic selection had more moderate binding scores, compared to EMID1, except for a small patch towards the end of THR. This particular peak falls within a region of negatively selected DRBS. **Right:** In NTN1, a modest number of DRBS covered the signal peptide and the NTR domain, not showing a particularly high binding scores. **F) Selection on COL1A1 hydroxyprolines**. Relatively strong negative selection was detected on the majority of sites, with only two residues, P587 and P919, under pervasive positive and episodic selection, respectively.

#### Disordered Binding Sites Prediction and Selection

Although UniProt already annotates disordered regions, we further analyzed the three proteins for disordered region binding sites (DRBS) at AIUPred to gain a better understanding of their functional potential. The proteins differed markedly in DRBS content: nearly half of EMID1 and over two-thirds of COL1A1 were spanned by DRBS, while they were almost absent in NTN1. Across all three molecules, the vast majority of DRBS sites were under some form of selection (Figure 5D). As expected, negative selection dominated, but a notable proportion of sites in EMID1 and COL1A1 showed episodic and/or much rarely pervasive positive selection. DRBS binding scores also varied, with the highest binding scores typically detected for sites under strong negative selection (Figure 5E). In EMID1, the C-terminal region contained high-scoring DRBS under both negative and episodic selection, while in COL1A1, episodic selection was associated with moderately high binding scores. Finally, the DRBS-rich THR domain in COL1A1 harbors numerous hydroxyprolines (Figure 6F, E), most of which are under relatively strong negative selection.

**Figure 6.**
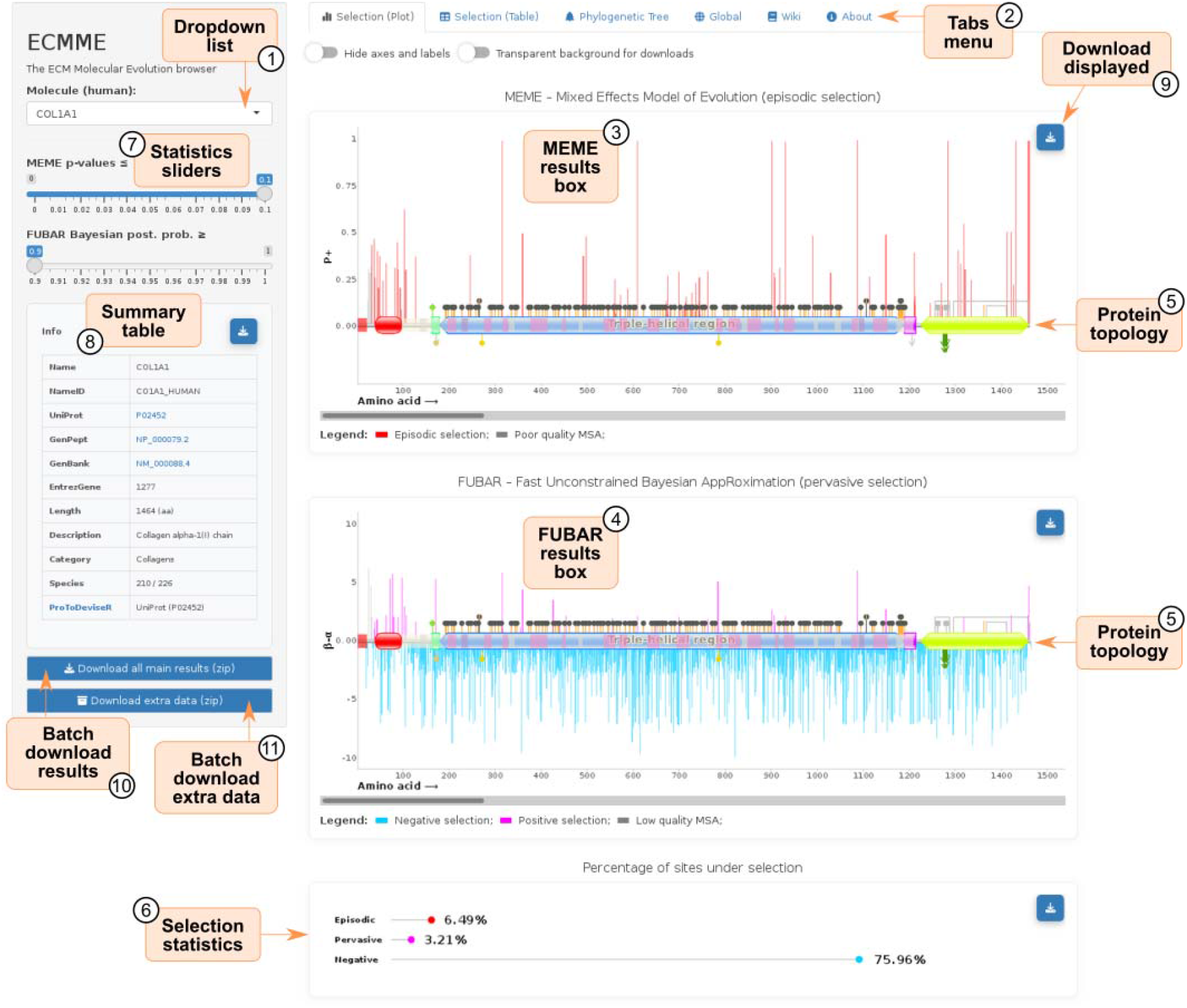
ECMME interface. The left sidebar contains a searchable dropdown menu listing all 272 core ECM proteins (1). Selecting a protein loads the data into the main panel, which opens by default on the “Selection (Plot)” tab (2). The results from MEME and FUBAR analyses are shown in dedicated boxes (3, 4) plotted onto the protein’s topology (5). A concise summary of the overall selection statistics across all sites is provided at the bottom (6). Users can interactively filter the displayed sites using sliders in the sidebar (7) to adjust thresholds for p-values (MEME) or Bayesian posterior probabilities (FUBAR). Immediately below the sliders, a summary table (8) displays metadata for the selected protein, including database identifiers (ID), the number of orthologous species included in the analysis, and the ID used to generate the topology diagram at ProToDeviseR. Each visualization component can be downloaded individually using its respective download button (9). In addition, the sidebar provides batch download buttons for all results of the selected protein (10), as well as a separate option (11) to download supplementary data files. The top tab menu (2) also gives access to a detailed table view of the selection results, the phylogenetic tree used for the analysis, global statistics for the entire dataset, and dedicated Wiki and About sections.

### ECMME, an open access database of molecular evolution within the ECM

Giving a detailed overview of each of the 272 molecules analyzed in this study goes beyond the scope of the paper, therefore, we created the ECM Molecular Evolution (ECMME) online library. We devised ECMME with an ease-to-use in mind, aiming to make browsing, searching and visualizing the data convenient, only a click away. The graphical user interface (GUI) is available online (https://izzilab-ecmme.share.connect.posit.cloud/) and requires no local installation or setting up by the users.

The GUI features a left side-bar, with a drop-down list of the core ECM molecules from human and statistics threshold sliders (Figure 6). The main panel contains four results-dedicated tabs: “Selection (Plot)”, “Selection (Table)”, “Phylogenetic tree”, “Global”, and additionaly a “Wiki” page and an “About” section.

Upon choosing a molecule from the drop-down list, a small sumamry table appears on the left, and the evolutionary selection results will be displayed graphically in the “Selection (Plot)” tab. Sites inferred under selection by MEME and FUBAR are plotted onto the protein topology of the protein from human, and basic sites selection statistics for the molecule are shown at the bottom. Episodic selection is shown in red, pervasive positive selection is reported in purple, while negative selection is denoted in light-blue. Selected sites from MSA columns considered as “unreliable” or difficult to align, are marked in gray, regardless of the selection type. This does not necessarily make results from such regions wrong, simply they should be considered with more care by the users. Sliders in the left side-bar allow for further filtering of the results based on p-values (0 - 0.1) for MEME and Bayes posterior probability for FUBAR (0.9 - 1). The selection data dynamically updates upon changing the sliders values, and in addition to the graphics, is offered as a table in the “Selection (Table)” tab. The “Phylogenetic tree” tab, displays the species used in the analysis of the chosen molecule, with the option to show them in the context of the global tree. The “Global” tab contains selection statistics covering all molecules, at the default cutoff thresholds of p-values and Bayes posterior probability for MEME and FUBAR. There, ECM components can be ranked by negative, positive pervasive and episodic selection, while a table overview is included underneath. Additionally, a chart ranking molecules by the number of species used for the analysis of each one is generated at the bottom. If a molecule is selected from the drop-down list on the left, it will be pointed in the charts and highlighted in the table. The “Wiki” and “About” tabs provide information on the results interpretation and ECMME itself.

Results are downloadable, either individually or in batch, allowing users to incorporate the ready charts in papers or presentations and, if desired, to run the analyses themselves.

## Discussion

We present a per-residue map of the natural selection pressures operating on the core matrisome of placental mammals, delivered through ECMME. This open and interactive resource allows for direct exploration at the level of proteins, domains, motifs, and residues.

Across the dataset, the dominant signal is strong purifying selection, consistent with the structural and functional indispensability of the extracellular matrix. This constraint, however, is not absolute: nearly every molecule also carries localized signatures of episodic and, more rarely, pervasive positive selection Collagens widely regarded as the most rigidly constrained ECM class, yet carry a significantly elevated levels of episodic positive selection, indicating that even canonical structural proteins retain pockets of adaptive flexibility. Certain structural compartments, such as, the peri-vascular ECM and basement membrane appear preferentially enriched for episodic selection relative to growth-factor–binding proteins, a trend not attributable to collagens alone. Together, these patterns suggest that architectural ECM niches are favorable substrates for species-specific adaptation against an otherwise conserved background.

At the single-molecule scale, mapping selection onto protein topology clearly illustrates the spatial, non-random, organization of these regimes. Typically, we observed strong overall conservation punctuated by isolated point or clustered adaptive changes. In our exemplar proteins, episodic selection in EMID1 concentrated at domain borders, pervasive positive selection in COL1A1 clustered around the VWFC domain, while the strongest purifying selection in NTN1 was within its NTR domain. Selection signals were structured across functional elements: short linear motifs were largely conserved, while specific cleavage and glycosaminoglycan-attachment motifs showed episodic diversification. Disordered binding sites combined strong purifying selection with a notable fraction of adaptively evolving sites. Even post-translational modification sites, such as COL1A1 triple-helical hydroxyprolines, were overwhelmingly under negative selection, underscoring their functional weight.

The core matrisome thus emerges not as an evolutionarily inert scaffold but rather as a dynamic system, where deep conservation and punctuated adaptive fine-tuning coexist within the same molecule. This reflects the long-recognized fundamental duality of the ECM as being both ancient and continually reshaped. We expect this resource to be widely useful for prioritizing functional residues for direct mutagenesis, interpreting disease variants, and projecting selection onto protein structures and binding interfaces.

By combining a transparent analysis pipeline with an accessible web browser and downloadable datasets, ECMME makes the results of this study easily available to researchers. Its user-friendly interface allows anyone to quickly search for a molecule of interest, interactively visualize selection patterns in real time, and explore the data in the context of protein domains, motifs, and structural features. By doing so, ECMME facilitates the integration of molecular evolution insights into broader ECM research and provides a well-documented framework that can be extended to wider the knowledge on matrisome and beyond.

## Data Availability

The data generated in this study and used by the ECMME RShiny app is available at the groups’s GitHub repository (https://github.com/Izzilab/ecmme/). Additional files referred to in this study are: Additional File 1 - Global_table.csv (UniProt/RefSeq IDs match, proteins metadata and a summary of the selection results for all genes), Additional File 2 - genes_species_matrix.csv (molecules and the species where a good quality sequence orthologue was found and used in the study), Additional File 3 - tree.nwk (global phylogenetic tree obtained from TimeTree).

Datasets used by ECMME (internally) are: allgenessel.rda (UniProt/RefSeq IDs match, proteins metadata and a summary of the selection results for all genes), fasta.codon.rda (codon alignments used by HyPhy), orthol.genesTrees.rda (phylogenetic trees used by HyPhy, trimmed from the global tree to represent the orthologues species of each gene), genes_species_matrix.csv (molecules and the species where a good quality sequence orthologue was found and used in the study), orthol_xm.rda (RefSeq identifiers for the orthologues of each gene), proteins.feature.table.rda (protein features as a table), proteins.json.rda (protein features in JSON format used by ProToDeviseR to generate topology schemes), seq.reference.rda (MSA quality, as determined by Gblocks plotted against the reference amino acid sequence from human), tree.rda (global phylogenetic tree obtained from TimeTree), trimmed.hyphy.rda (results from MEME and FUBAR, filtered by p-value < 0.1 and Bayes PP > 0.9).

Protein topology schemes are available in the www/images/subfolder. Non-filtered HyPhy output data is deposited on Zenodo (doi: 10.5281/zenodo.20592814): json.results.rda (MEME and FUBAR results in JSON format).

## Code Availability

The code of ECMME RShiny app is available at https://github.com/Izzilab/ecmme/. The code of the data preparation pipeline is available at https://github.com/Izzilab/ecmme-pipeline/. Development was done on CRUX 3.8 (https://crux.nu/) distribution of GNU/Linux with all necessary software installed from the ports at the distribution’s port database (https://crux.nu/portdb/), where ports for the R packages ecosystem were generated by cran2crux [15].

## Methods

ECMME is written in R (v4.5, https://www.r-project.org/) and its source code is freely available at our GitHub repository (https://github.com/izzilab/ecmme) under GPLv3 license. The graphical user interface (GUI) is implemented in R Shiny to offer dynamic results browsing and display.

### ECM evolutionary selection pipeline

The preparation and results processing pipeline is also designed in R, with several wrappers written in BASH, to drive external programs and handle downloads. As with ECMME, the code is available on our GitHub repository (https://github.com/Izzilab/ecmme-pipeline/), under GPLv3 license. All database (DB) and knowledge-base (KB) accessions were done on 08 January 2026. R libraries rentrez, seqinr and ape, were used for DB accession, sequence work and phylogenetic tree processing, respectively. An brief overview of the procedure is given in the sections below.

### ECM core proteins canonical isoforms and homologues retrieval

The Matrisome project [13] list was used to subset the 274 core members (*Homo sapiens*) from the full list. They were cross-referenced to GeneCards [16] by their *entrez_id*, in order to retrieve the respective up-to-date UniProtKB identifiers. Protein information (*.txt format) was downloaded for each entry from UniProt, and the “Displayed” UniProt IDs and their corresponding RefSeq IDs were extracted.

Amino acid (aa) sequences were then downloaded from UniProt and corresponding coding DNA sequences were downloaded (CDS) from RefSeq. CDS were translated and compared to the aa sequences from UniProt, to set aside any discrepancies between the databases. If a mismatch was found, the latest RefSeq sequence was retrieved and compared again. However, in the case there was no consensus between the databases, the RefSeq CDS ID listed by GeneCards was considered, instead. Finally, the corresponding CDS were collected and designated hereafter as the “reference” sequences from *Homo sapiens*.

The NCBI DataSets [17] package (v18.14.0) was used to retrieve all sequences at level *placentalia* for each gene, by their *entrez_id*.

### Species and a phylogenetic tree

Collectively, the retrieved homologues covered initially a total of 234 placental mammals. The list of species was submitted to the TimeTree [14] KB, species not found in the KB were removed, while alternative species names were considered where necessary. We prepared a final, manually curated, list of 228 species (Supplementary Table S1, Figure S1), which was resubmitted to TimeTree and the phylogenetic tree was downloaded in *.nwk format (Additional File 3).

### Homologues sequences quality evaluation

A series of quality checks were performed on the downloaded CDS, by inspecting for non-standard nucleotides, non-standard amino acids in the translated sequences, and in-frame stop codons. Such entries were discarded, including, sequences from species not found at TimeTree. Finally, the CDS that passed the quality criteria had their terminal stop codons removed and all sequences were translated, in preparation for the next steps.

### Identification of closest orthologues to the reference sequence

A reciprocal BLASTP [18] was carried out per gene entry locally, for the isoforms from each species against the reference sequence from human, using default settings. After the run, BLAST hits were filtered by the following criteria: E-value (< 1e-5), percentage of identical matches cut-off (> 75), query (Q) / subject (S) minimum and maximum length ratio (> 0.1 <), maximum absolute start of the Q and S alignments difference (< 35 aa), maximum relative start of the Q and S alignments difference (< 0.1), Q coverage as percent (> 40). A composite score was calculated by multiplying the bit-score reported in the results by the percentage of identical matches. The composite score was used to rank the BLAST hits, following the initial filtering. The highest ranking sequence from each species was then collected, per gene, together with the reference sequence from human.

We excluded SSPOP, a pseudogene, and a POMZP3, for which only the human sequence passed our filtering criteria, leaving 272 molecules (Additional File 1).

### Multiple sequence alignments

The collected orthologous sequences for each gene entry were aligned by MUSCLE5 [19] and their corresponding CDS were then codon-aligned by pal2nal [20]. The quality of the protein multiple sequence alignments (MSA) was evaluated by Gblocks [21], for later referral.

### Evolutionary selection screening

Two methods from the HyPhy suite (v2.5.93), MEME [22] and FUBAR [23], were used to test for evolutionary selection on each gene. The codon alignments, and trees pruned from the global tree in line with the orthologues of each gene, were submitted to the program.

### Post-run processing

The results files (*.json format) produced by HyPhy were processed and filtered by the default MEME and FUBAR thresholds: p-value < 0.1 and Bayes post. prob. > 0.9, respectively. The reference amino acid sequence from *Homo sapiens* was added alongside its respective coordinates in the MSA. The quality of the MSA at each amino acid position was also appended.

Protein topology schemes were first batch generated (*.json format) by ProToDeviseR [24]. Files were submitted to a slightly modified graphics generator (currently in the *testing* branch at the tool’s GitHub) to render the schemes graphically. The selected sites, as determined by MEME and FUBAR, were plotted onto the protein topology of the amino acid sequence of each gene.

Additionally, functions were created to display the phylogenetic tree, rank the genes by selection, by number of species used in the HyPhy runs, and more.

### Further sequence characterization

Short functional motifs (SFM) were predicted at the Eukaryotic Linear Motif resource [25], setting cell compartment to “extracellular”. Scanning for disordered region binding sites was performed at AIUPred [26,27] and results were filtered for both disordered and binding values to be over 0.5.

### Data visualization

Plots were done in R and Gnumeric (https://gitlab.gnome.org/GNOME/gnumeric), Venn diagrams were generated by DeepVenn [28], scalable vector graphics (SVG) of animals were from SVG REPO (https://staging.svgrepo.com/) and figures were assembled in Inkscape (https://inkscape.org/).

## Author Contributions

Conceptualization: VI, PBP; coding: PBP; data analysis: PBP, VI; figures design: PBP; code testing and assessment: AO, VI, PBP; funding acquisition: VI, JR; manuscript preparation: PBP, VI; project supervision: VI; manuscript review and editing: all authors.

