## Supplementary for "ECMME: an atlas of selection pressures on the mammalian extracellular matrix reveals contrasting evolutionary dynamics"

**Table S1. Latin and common name of the 228 species used.** Order follows the phylogenetic tree in Fig 2 and Fig S1.

| Latin name | Common name(s) |
|---|---|
| *Mirounga angustirostris* | Northern elephant seal |
| *Mirounga leonina* | Southern elephant seal |
| *Leptonychotes weddellii* | Weddell seal |
| *Neomonachus schauinslandi* | Hawaiian monk seal |
| *Phoca vitulina* | Harbor seal |
| *Halichoerus grypus* | Gray seal |
| *Zalophus californianus* | California sea lion |
| *Eumetopias jubatus* | Steller sea lion |
| *Callorhinus ursinus* | Northern fur seal |
| *Odobenus rosmarus divergens* | Pacific walrus |
| *Mustela lutreola* | European mink |
| *Mustela putorius furo* | Domestic ferret |
| *Mustela nigripes* | Black-footed ferret |
| *Mustela erminea* | Stoat |
| *Neogale vison* | American mink |
| *Enhydra lutris kenyoni* | Northern sea otter |
| *Lutra lutra* | Eurasian otter |
| *Lontra canadensis* | North American river otter |
| *Meles meles* | European badger |
| *Ursus maritimus* | Polar bear |
| *Ursus arctos* | Brown bear |
| *Ursus americanus* | American black bear |
| *Ailuropoda melanoleuca* | Giant panda |
| *Vulpes lagopus* | Arctic fox |
| *Vulpes vulpes* | Red fox |
| *Nyctereutes procyonoides* | Raccoon dog |
| *Canis aureus* | Golden jackal |
| *Canis lupus familiaris* | Domestic dog |
| *Herpailurus yagouaroundi* | Jaguarundi |
| *Puma concolor* | Puma |
| *Acinonyx jubatus* | Cheetah |
| *Lynx canadensis* | Canada lynx |
| *Lynx rufus* | Bobcat |
| *Prionailurus viverrinus* | Fishing cat |
| *Prionailurus bengalensis* | Leopard cat |
| *Felis catus* | Domestic cat |
| *Leopardus geoffroyi* | Geoffroy's cat |
| *Panthera pardus* | Leopard |
| *Panthera leo* | Lion |
| *Panthera onca* | Jaguar |
| *Panthera uncia* | Snow leopard |
| *Panthera tigris* | Tiger |
| *Neofelis nebulosa* | Clouded leopard |
| *Crocuta crocuta* | Spotted hyena |
| *Hyaena hyaena* | Striped hyena |
| *Suricata suricatta* | Meerkat |
| *Manis pentadactyla* | Chinese pangolin |
| *Manis javanica* | Sunda pangolin |

| | |
|---|---|
| *Equus quagga* | Plains zebra |
| *Equus asinus* | Donkey |
| *Equus caballus* | Horse |
| *Equus przewalskii* | Przewalski's horse |
| *Diceros bicornis minor* | Southern black rhinoceros |
| *Ceratotherium simum simum* | Southern white rhinoceros |
| *Tursiops truncatus* | Common bottlenose dolphin |
| *Delphinus delphis* | Short-beaked common dolphin |
| *Pseudorca crassidens* | False killer whale |
| *Globicephala melas* | Long-finned pilot whale |
| *Sagmatias obliquidens* | Pacific white-sided dolphin |
| *Lagenorhynchus albirostris* | White-beaked dolphin |
| *Orcinus orca* | Killer whale |
| *Phocoena phocoena* | Harbor porpoise |
| *Phocoena sinus* | Vaquita |
| *Neophocaena asiaeorientalis asiaeorientalis* | Yangtze finless porpoise |
| *Monodon monoceros* | Narwhal |
| *Delphinapterus leucas* | Beluga whale |
| *Lipotes vexillifer* | Baiji |
| *Mesoplodon densirostris* | Blainville's beaked whale |
| *Physeter macrocephalus* | Sperm whale |
| *Kogia breviceps* | Pygmy sperm whale |
| *Balaenoptera musculus* | Blue whale |
| *Balaenoptera ricei* | Rice's whale |
| *Eschrichtius robustus* | Gray whale |
| *Balaenoptera acutorostrata* | Common minke whale |
| *Eubalaena glacialis* | North Atlantic right whale |
| *Hippopotamus amphibius kiboko* | Eastern hippopotamus |
| *Ovis aries* | Domestic sheep |
| *Ovis canadensis* | Bighorn sheep |
| *Capra hircus* | Domestic goat |
| *Budorcas taxicolor* | Takin |
| *Capricornis sumatraensis* | Sumatran serow |
| *Oryx dammah* | Scimitar-horned oryx |
| *Bos indicus* | Zebu |
| *Bos taurus* | Domestic cattle |
| *Bison bison bison* | American bison |
| *Bos mutus* | Wild yak |
| *Bubalus bubalis* | Water buffalo |
| *Moschus berezovskii* | Forest musk deer |
| *Cervus elaphus* | Red deer |
| *Cervus canadensis* | Elk |
| *Dama dama* | Fallow deer |
| *Muntiacus reevesi* | Reeves's muntjac |
| *Odocoileus virginianus* | White-tailed deer |
| *Sus scrofa* | Wild boar |
| *Phacochoerus africanus* | Common warthog |
| *Camelus bactrianus* | Bactrian camel |
| *Camelus ferus* | Wild Bactrian camel |
| *Camelus dromedarius* | Dromedary |
| *Vicugna pacos* | Alpaca |

| | |
|---|---|
| *Myotis myotis* | Greater mouse-eared bat |
| *Myotis daubentonii* | Daubenton's bat |
| *Myotis davidii* | David's myotis |
| *Myotis yumanensis* | Yuma myotis |
| *Myotis brandtii* | Brandt's bat |
| *Myotis lucifugus* | Little brown bat |
| *Eptesicus fuscus* | Big brown bat |
| *Pipistrellus kuhlii* | Kuhl's pipistrelle |
| *Miniopterus natalensis* | Natal long-fingered bat |
| *Molossus molossus* | Pallas's mastiff bat |
| *Phyllostomus discolor* | Pale spear-nosed bat |
| *Phyllostomus hastatus* | Greater spear-nosed bat |
| *Artibeus jamaicensis* | Jamaican fruit bat |
| *Sturnira hondurensis* | Honduran yellow-shouldered bat |
| *Desmodus rotundus* | Common vampire bat |
| *Pteronotus mesoamericanus* | Mesoamerican mustached bat |
| *Saccopteryx bilineata* | Greater sac-winged bat |
| *Saccopteryx leptura* | Lesser sac-winged bat |
| *Pteropus vampyrus* | Large flying fox |
| *Pteropus medius* | Indian flying fox |
| *Pteropus alecto* | Black flying fox |
| *Rousettus aegyptiacus* | Egyptian rousette |
| *Rhinolophus ferrumequinum* | Greater horseshoe bat |
| *Rhinolophus sinicus* | Chinese horseshoe bat |
| *Hipposideros armiger* | Great Himalayan leaf-nosed bat |
| *Sorex araneus* | Common shrew |
| *Sorex fumeus* | Smoky shrew |
| *Suncus etruscus* | Etruscan shrew |
| *Erinaceus europaeus* | European hedgehog |
| *Talpa occidentalis* | Western mole |
| *Condylura cristata* | Star-nosed mole |
| *Microtus oregoni* | Creeping vole |
| *Microtus pennsylvanicus* | Meadow vole |
| *Microtus ochrogaster* | Prairie vole |
| *Alexandromys fortis* | Reed vole |
| *Chionomys nivalis* | European snow vole |
| *Arvicola amphibius* | Eurasian water vole |
| *Myodes glareolus* | Bank vole |
| *Cricetulus griseus* | Chinese hamster |
| *Mesocricetus auratus* | Syrian hamster |
| *Phodopus roborovskii* | Roborovski dwarf hamster |
| *Peromyscus eremicus* | Cactus mouse |
| *Peromyscus californicus insignis* | California mouse |
| *Peromyscus leucopus* | White-footed mouse |
| *Peromyscus maniculatus bairdii* | Deer mouse |
| *Onychomys torridus* | Southern grasshopper mouse |
| *Mus caroli* | Ryukyu mouse |
| *Mus musculus* | House mouse |
| *Mus pahari* | Southeast Asian house mouse |
| *Mastomys coucha* | Southern multimammate mouse |
| *Apodemus sylvaticus* | Wood mouse |

| | |
|---|---|
| *Arvicanthis niloticus* | Nile grass rat |
| *Grammomys surdaster* | African woodland thicket rat |
| *Rattus norvegicus* | Norway rat |
| *Meriones unguiculatus* | Mongolian gerbil |
| *Psammomys obesus* | Fat sand rat |
| *Acomys russatus* | Golden spiny mouse |
| *Nannospalax galili* | Upper Galilee blind mole-rat |
| *Jaculus jaculus* | Lesser Egyptian jerboa |
| *Dipodomys ordii* | Ord's kangaroo rat |
| *Dipodomys merriami* | Merriam's kangaroo rat |
| *Dipodomys spectabilis* | Banner-tailed kangaroo rat |
| *Perognathus longimembris pacificus* | Pacific pocket mouse |
| *Castor canadensis* | North American beaver |
| *Chinchilla lanigera* | Long-tailed chinchilla |
| *Octodon degus* | Common degu |
| *Cavia porcellus* | Domestic guinea pig |
| *Fukomys damarensis* | Damaraland mole-rat |
| *Heterocephalus glaber* | Naked mole-rat |
| *Marmota monax* | Groundhog |
| *Marmota marmota marmota* | Alpine marmot |
| *Marmota flaviventris* | Yellow-bellied marmot |
| *Callospermophilus lateralis* | Golden-mantled ground squirrel |
| *Urocitellus parryii* | Arctic ground squirrel |
| *Ictidomys tridecemlineatus* | Thirteen-lined ground squirrel |
| *Sciurus carolinensis* | Eastern gray squirrel |
| *Ochotona curzoniae* | Plateau pika |
| *Ochotona princeps* | American pika |
| *Lepus europaeus* | European hare |
| *Oryctolagus cuniculus* | European rabbit |
| *Macaca fascicularis* | Crab-eating macaque |
| *Macaca mulatta* | Rhesus macaque |
| *Macaca thibetana thibetana* | Tibetan macaque |
| *Macaca nemestrina* | Southern pig-tailed macaque |
| *Cercocebus atys* | Sooty mangabey |
| *Mandrillus leucophaeus* | Drill |
| *Theropithecus gelada* | Gelada |
| *Papio anubis* | Olive baboon |
| *Chlorocebus sabaeus* | Green monkey |
| *Rhinopithecus bieti* | Black snub-nosed monkey |
| *Rhinopithecus roxellana* | Golden snub-nosed monkey |
| *Trachypithecus francoisi* | François' langur |
| *Colobus angolensis palliatus* | Angolan colobus |
| *Piliocolobus tephrosceles* | Ugandan red colobus |
| *Pan troglodytes* | Common chimpanzee |
| *Pan paniscus* | Bonobo |
| *Homo sapiens* | Human |
| *Gorilla gorilla gorilla* | Western gorilla |
| *Pongo abelii* | Sumatran orangutan |
| *Pongo pygmaeus* | Bornean orangutan |
| *Hylobates moloch* | Silvery gibbon |
| *Symphalangus syndactylus* | Siamang |

| | |
|---|---|
| *Nomascus leucogenys* | Northern white-cheeked gibbon |
| *Sapajus apella* | Tufted capuchin |
| *Cebus imitator* | White-faced capuchin |
| *Saimiri boliviensis* | Bolivian squirrel monkey |
| *Callithrix jacchus* | Common marmoset |
| *Aotus nancymaae* | Nancy Ma's night monkey |
| *Carlito syrichta* | Philippine tarsier |
| *Eulemur rufifrons* | Red-fronted lemur |
| *Lemur catta* | Ring-tailed lemur |
| *Microcebus murinus* | Gray mouse lemur |
| *Propithecus coquereli* | Coquerel's sifaka |
| *Otolemur garnettii* | Northern greater galago |
| *Nycticebus coucang* | Sunda slow loris |
| *Galeopterus variegatus* | Sunda flying lemur |
| *Cynocephalus volans* | Philippine flying lemur |
| *Tupaia chinensis* | Chinese tree shrew |
| *Tenrec ecaudatus* | Tailless tenrec |
| *Echinops telfairi* | Lesser hedgehog tenrec |
| *Chrysochloris asiatica* | Cape golden mole |
| *Elephantulus edwardii* | Cape rock elephant shrew |
| *Orycteropus afer afer* | Aardvark |
| *Elephas maximus indicus* | Indian elephant |
| *Loxodonta africana* | African savanna elephant |
| *Trichechus manatus latirostris* | Florida manatee |
| *Choloepus didactylus* | Southern two-toed sloth |
| *Tamandua tetradactyla* | Southern tamandua |
| *Dasypus novemcinctus* | Nine-banded armadillo |

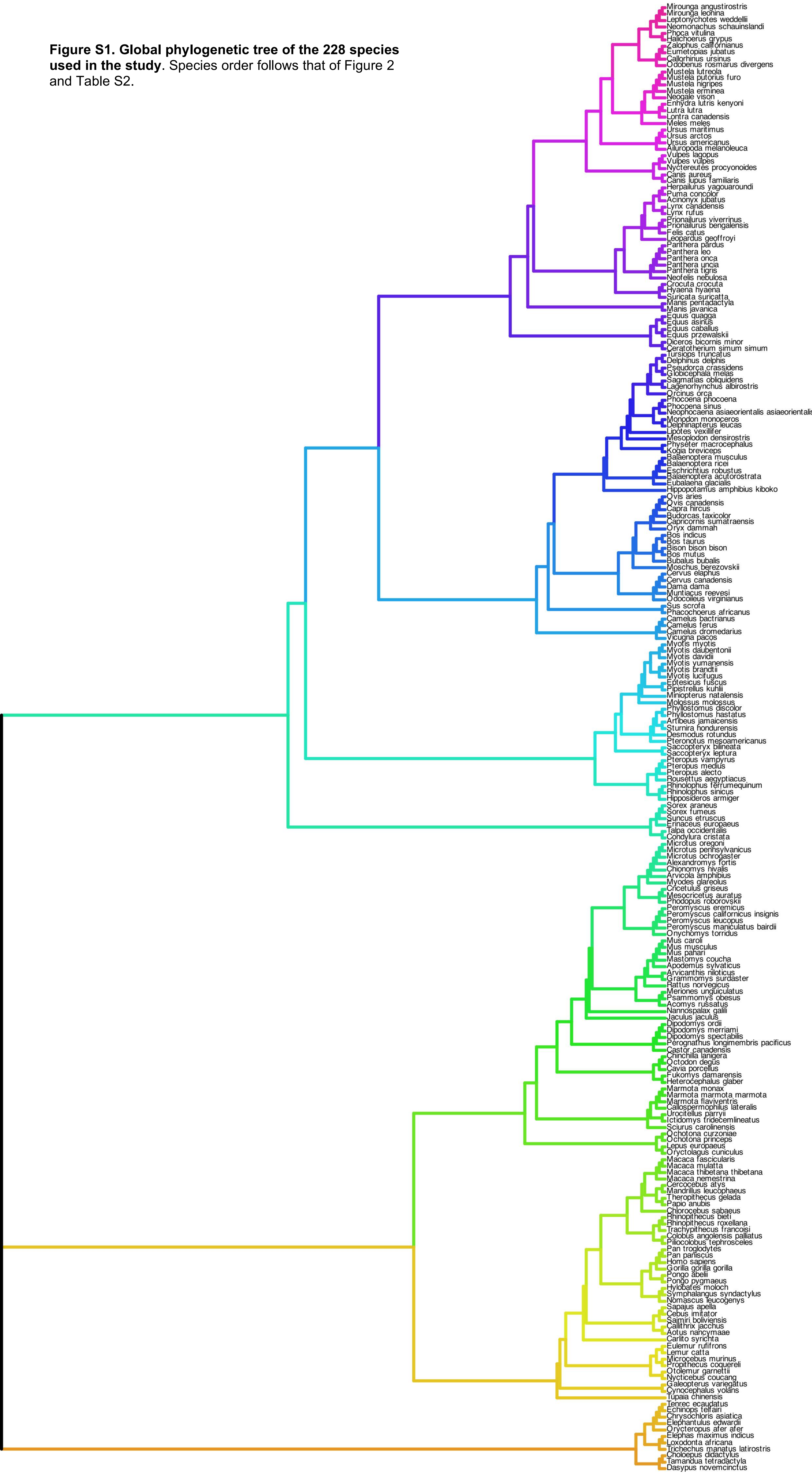


**Figure S1. Global phylogenetic tree of the 228 species used in the study**. Species order follows that of Figure 2 and Table S2.
